# Long-Term Colonization Dynamics of Probiotic *Aliivibrio* spp. in Atlantic Salmon (*Salmo salar) Following Bath Administration*

**DOI:** 10.1101/2024.12.21.629900

**Authors:** Marius Steen Dobloug, Stanislav Iakhno, Simen Foyn Nørstebø, Øystein Evensen, Henning Sørum

## Abstract

Ulcerative conditions present a major challenge in Norwegian salmon farming. Probiotic Aliivibrio species have previously been demonstrated to provide health benefits in both Atlantic salmon and lumpfish, although the underlying mechanisms and the host-bacteria interactions remain unclear. This study aimed to investigate whether these bacteria could colonize Atlantic salmon following bath administration, determine the tissue tropism, and assess the duration of colonization. We examined the host microbiota using culture-based methods, qPCR and immunohistochemistry techniques specifically designed to target the applied Aliivibrio strains. Our findings reveal that the probiotic bacteria can successfully colonize Atlantic salmon and persist for at least nine months post-administration. We identified the administered strains in the skin and underlying tissue with all three methods. The probiotics were also identified in the distal intestine and the visceral organs. Additionally, we isolated the probiotic Aliivibrio species from mixed cultures in ulcerated areas. While viable bacteria were recoverable from recently euthanized fish, tissue decay promoted bacterial recovery of the administered species across all experiments. Given prior evidence on ulcer reduction associated with these probiotics, competitive exclusion appears to be a plausible mechanism of action, though further investigation is warranted.

## 1. Introduction

The global human population is rising, making fisheries and aquaculture essential for global food security. For the last two decades, the increased production in this sector has primarily come from aquaculture, as wild-caught fish are naturally limited ^1^. According to the Food and Agriculture Organization (FAO), Norway accounted for 45 % of all aquaculture production in Europe in 2020 and is currently ranked as the world’s second-largest producer of marine and coastal fin fish. Atlantic salmon is the primary farmed species in Norway, with its annual production in million tonnes tripling over the past two decades ^2^.

Rapid industry growth stagnated in 2012 due to biological and environmental challenges. These challenges remain unresolved, leading to unprecedented mortalities in 2023, surpassing 100 million fish dying before reaching slaughter ^3,4^. Since the 1990s, vaccines have limited the prevalence of acute bacterial diseases in Norwegian aquaculture ^5^. However, vaccines show limited effectiveness towards Moritella viscosa, which is one of the main causes of the recent increase in ulcer development and mortality ^4^. Antibiotic use to control bacterial infections in salmon aquaculture is limited by self-directed restriction and government policies due to the global threat of antibiotic resistance development, leaving the aquaculture industry vulnerable ^6^. The current situation opens the door for alternative measures for disease control.

Probiotic bacteria are live microorganisms that confer a health benefit to the host when administered in adequate amounts ^7,8^. Probiotic bacteria may suppress pathogens by competitive exclusion with modes of actions such as blocking of attachment sites on mucosal surfaces ^9–12^, antagonistic activity due to the production of antimicrobial compounds ^13,14^ ^14,15^, or by seizing key resources such as iron ^16–21^. Probiotic bacteria may also support the host in other ways, such as through immunomodulation ^21–25^ or enzyme production improving host digestion ^26^. While the use of antibiotics remains a common practice in aquaculture worldwide, it has been advocated for a transition to more sustainable alternatives, such as probiotics ^27^. However, it has been argued that the use of probiotics is currently limited due to a lack of awareness among farmers regarding their value and application ^27^. The success of probiotics depends on the specific strain used and the route of administration. Numerous potential probiotics for fish have been explored, with lactic acid bacteria or Bacillus species being the most commonly utilized, typically administered by feed ^27–30^. A novel probiotic approach for Atlantic salmon was previously demonstrated, utilizing a combination of Aliivibrio species administered through the rearing water.

The Vibrionaceae family contains many diverse aquatic bacteria, including the Aliivibrio genus ^31^. There are currently six established species within this genus: Aliivibrio logei, Aliivibrio salmonicida, Aliivibrio wodanis, Aliivibrio sifiae, Aliivibrio finisterrensis, and Aliivibrio fischeri ^32^. All six species have previously been found in fish by culture or DNA-based methods. A. wodanis is typically associated with ulcers but has been detected on the intact skin of Atlantic salmon, as has A. logei and A. salmonicida ^33,34^. A. salmonicida, A. finisterrensis, A. logei, and A. wodanis have been isolated from the intestine of Atlantic salmon, while A. sifiae has been detected in the intestine of several other fish species ^35–39^. Uncharacterized Aliivibrio spp. have also been identified in the middle and distal intestine, head kidney, healthy skin and ulcers of Atlantic salmon ^39–41^. The luminescent A. fischeri is one of the most well-studied symbiotic bacteria housed within the light organ of aquatic species such as Monocentris japonica and Euprymna scolopes ^42,43^. In general, the Aliivibrio species are considered closely associated with fish, including Atlantic salmon.

Isolation of viable bacteria relies on culture-based techniques; however, this approach is challenging because only a small fraction of bacterial species is considered culturable ^44^. The discrepancy between viable plate counts and direct microscopic counts is commonly referred to as “The great plate count anomaly” ^45^. Most bacteria found in humans ^46,47^ and teleosts are also regarded as non-culturable ^48,49^. Some culturable bacteria can also transition into non-culturable states known as VBNC ^50^. Colwell et al. introduced the term “viable but nonculturable (VBNC)” in 1985, and VBNC has since been defined by James D. Oliver as “a cell which can be demonstrated to be metabolically active, while being incapable of undergoing a sustained cellular division required for growth in or on a medium normally supporting growth of that cell” ^51,52^. This survival state was initially described for Vibrio cholerae and a few other pathogens, but has since been reported for 115 species, many within the Vibrionaceae family ^53–56^. This includes A. fischeri ^57^, and one can assume that these states may occur in other species within the Aliivibrio genus. As interest in the VBNC state has grown, so have strategies for bringing bacteria back to culturable states ^50^. VBNC bacteria can be resuscitated in several ways, such as by specific stimulating factors or nutrients ^58–62^. Conditions for resuscitation can be provided by the host, evident by co-culture resuscitation in vitro using eukaryotic cell models ^63,64^, or in vivo using host models ^65–68^.

Homeostasis is how animals maintain a stable internal environment despite external changes. The immune system plays a vital role in homeostasis by protecting the body from disruptions. When an animal dies, the immune system no longer regulates the microbiota. Simultaneously, chemical alterations and autolysis disrupt cell membranes, releasing a vast reservoir of nutrients that enables bacterial proliferation and spread ^69^. Although bacteria may be non-culturable in the living host ^44^, post-mortem conditions can promote bacterial growth, enabling their re-isolation from the carcass. Decomposition is one of the most fundamental processes on Earth, as dead biological material is continuously recycled ^70^. The community of microorganisms driving the corpse decomposition and its necromass is known as the necrobiome, which is both dynamic and complex ^71^. This process is characterized by temporal shifts in bacterial populations, occurring in succession post-mortem, often called the “microbial clock” in forensic medicine ^72^. Despite the well-documented diversity of putrefaction-associated microbes, the human necrobiome has been shown to encompass conserved microbial networks with consistent community succession patterns ^70^. The necrobiome of fish is less studied, but the abundance of bacteria has been shown to increase by 250 times with significant changes to the composition in just 48 hours post-mortem ^73^. Fish have also been revealed to have well defined microbial community succession patterns independent of the water microbiota ^74^.

In recent work, it has been shown that the administration of two probiotic Aliivibrio species in Atlantic salmon is associated with improved survival rates, enhanced growth, a reduced prevalence of ulcers during M. viscosa outbreaks, and decreased salmon lice attachment ^75–77^. However, the mechanisms underlying these effects remain unknown. Since many key probiotic functions depend on the bacteria’s presence within the host, this study aims to determine whether these Aliivibrio species can colonize Atlantic salmon and evaluate the duration of colonization. We hypothesize that the administered bacteria can colonize the skin and the intestine, and tissue decay and temperature may influence their detection. By examining the salmon microbiota for the presence of these probiotic bacteria, we aim to improve the understanding of previously observed effects, potentially providing insights into the bacterial dynamics related to colonization, and to optimize this preventative approach in the health management of Atlantic salmon.

## 2. Materials and Methods

### 2.1 Fish and facilities

Male and female Atlantic salmon presmolt were sourced from Sørsmolt (trial 1, N = 45 fish; pilot, N = 5) and Fister Smolt (trial 2, N = 38 fish) and transferred by car to the Marine Research Station of the Norwegian Institute of Water Research (NIVA), Drøbak, Norway. Upon arrival, fish were placed in a holding tank containing 1000 litres of freshwater until probiotic administration. Fish welfare and water quality were surveyed, and care was provided by a responsible veterinarian or fish health biologist throughout the trial. Welfare issues resulting in death or euthanasia before the study endpoint were determined to be an exclusion criterion in advance of the study. No fish were excluded in either trial. The ARRIVE guidelines for the use of research animals have been followed. The Animal Welfare Approval (FOTS) numbers are 29937 and 30556.

All tanks in the NIVA research facility employ a flow-through system. All water is treated with UV irradiation (2 BetaLine-Eco BLE3.250L2/NW100/Us1/Mc/230V50Hz/PS, 100 m^3^/h maximum flow, UV T10 mm 95–98%, 45 mJ/cm^2^ UV dose) before entry to the research facility and emptied in a common collecting pipe; no water is shared between tanks. Freshwater is supplied from the same groundwater sand sedimental source, and all seawater is supplied from 50-meter depth in the Oslofjord. In trial 1 and the pilot, fish were kept in freshwater for two weeks after arrival before photoperiod smoltification of four weeks of 12:12 hours of light:darkness per day and two weeks of 24 hours of light per day. After smoltification, the fish were kept in saltwater of ≈33 ‰ salinity with a monthly average water temperature of 10 (±2.5) °C for the duration of the trial. In trial 2, the fish were kept in freshwater with ≈20 % seawater added to 7.4 (±0.9) ‰ salinity due to limitations of freshwater supply. The average water temperature in trial 2 was 10.4 (±1,2) °C.

### 2.2 Bacterial culture and probiotic bath administration

A combination of two Aliivibrio spp. strains were employed. Aliivibrio sp. strain Vl1, NCIMB 42593 and Aliivibrio sp. strain Vl2, NCIMB 42592 were originally isolated, cultivated and stored as detailed by Klakegg et al. ^75^. Both strains were collected from freeze stock at Previwo AS (Oslo, Norway) and subcultured to manufacture the probiotic product as explained by Klakegg et al. Harvested cultures were sampled for viable cell count and determination of purity by plating of serial dilutions and OD_600_ measurements. Samples were also collected from the probiotic bath, with effective doses determined by serial dilution plating to be ≈10^8^ colony forming units (CFU) per ml in both trials.

Probiotic bacteria were administered by immersing the fish for two minutes in a bath with probiotic bacteria diluted 1:20 in rearing water. The immersion bath contained 0.5 L probiotic broth culture and 19.5 L rearing water with 10‰ salinity and the anaesthetic chemical Benzoak Vet (ACD Pharmaceuticals AS, Leknes, Norway). The fish were then netted out and randomly distributed to holding tanks. Holding tanks for each group were randomly assigned based on a random number generator.

In trial 1, 30 fish were immersed in probiotics and 10 fish were mock immersed (no bacteria added to the bath). They were then weighed (70 grams on average) and separated into two different 180 L tanks, one for the probiotic immersed and one for the mock immersed, to let residual bacteria wash off. After 24 hours, they were pooled in the same 1000 L holding tank. In addition, five mock immersed control fish were kept in a separate 1000 L tank for the duration of the trial. In trial 2, three “absolute control fish” were euthanized upon arrival at the research facility to ensure they had not been in exposed to saltwater or the probiotic bacteria. The remaining 35 fish were weighed (56 grams on average) and distributed between 13 tanks, 10 for immersed fish and 3 for control fish. An overview of administration, incubation, and sampling in both trials of this study is provided in Figure 1.

**Figure 1:**
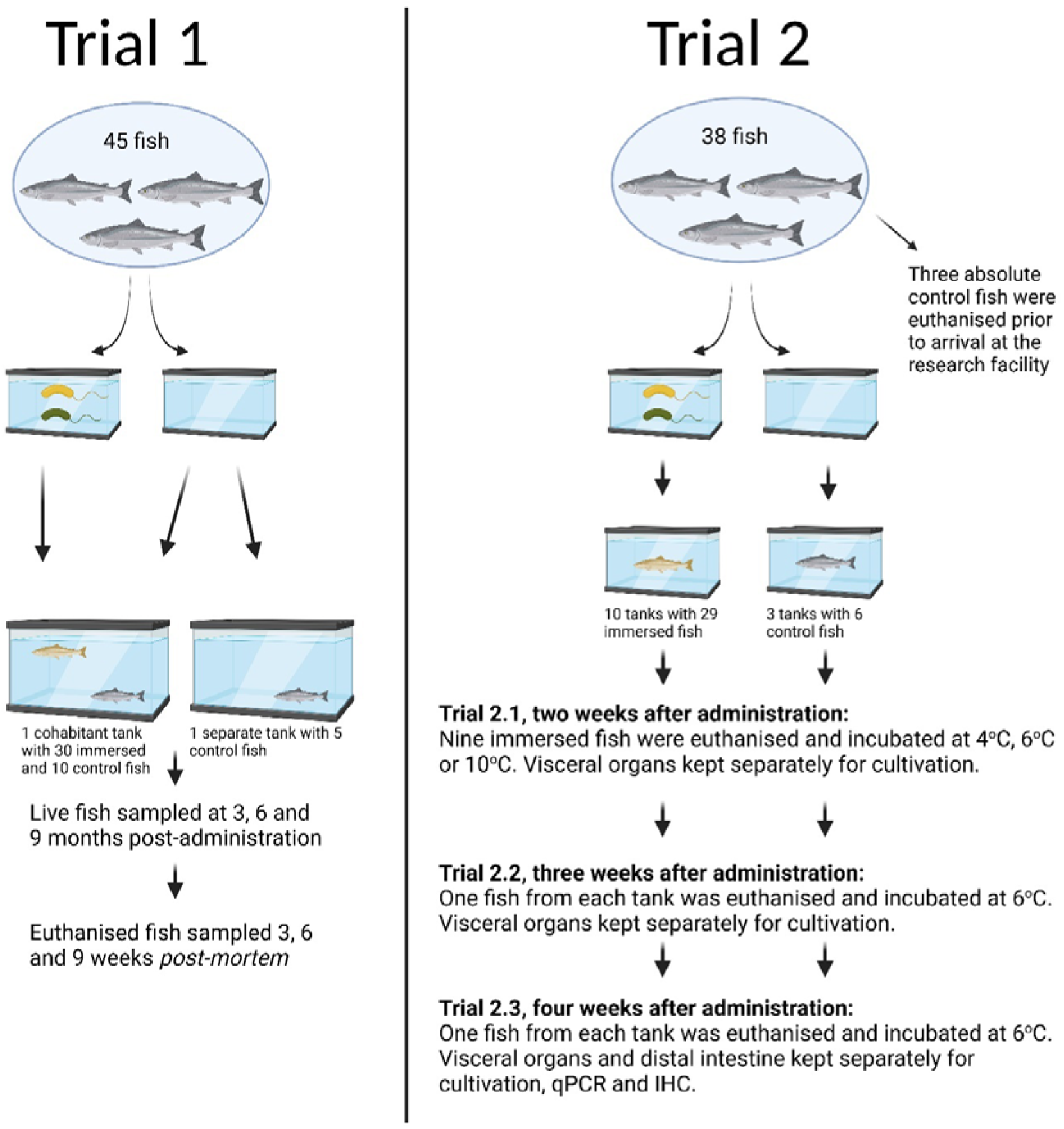
Overview of trial 1 and 2 (created with Biorender.com).

### 2.3 Incubation

In trial 1, nine months after bacterial bath administration, 30 immersed fish, 10 cohabitated control fish and five separate control fish (≈750 grams) were euthanized and individually placed into autoclaved polypropylene bags (Sarstedt, Germany). All 45 bags were incubated in a cool storage room of 6 (±2)°C for nine weeks.

The absolute control fish euthanized upon arrival at the research facility was incubated as in trial 1. In trial 2.1, two weeks after bacterial bath administration, nine immersed fish were euthanized. All fish were carefully dissected; an incision was made along the ventral side of the fish, and the visceral organs of all fish were removed and divided among three bags, each containing the pooled visceral organs of three fish. The remaining fish carcasses were placed in individual bags. To investigate if temperature affects bacterial recovery, the 12 bags containing fish or fish organs were incubated at 4 °C, 6 °C, or 10 °C, with 4 bags per temperature.

In trial 2.2, three weeks after bacterial bath administration, one fish from each tank (10 treated and 3 controls) was euthanized. All fish were dissected as previously described, but all samples (carcasses and visceral organs) were placed in individual sterile bags for 26 bags incubated at 6 °C.

In trial 2.3, four weeks after bacterial bath administration, the remaining fish from each tank (10 treated and 3 controls) were euthanized. All fish were dissected as previously described, but in addition to the carcasses and visceral organs, the distal intestine was also separated from the remaining visceral organs and placed in a separate sterile bag. This resulted in 39 bags which were all incubated at 6 °C.

### 2.4 Sampling

In trial 1, sampling was conducted by swabbing the skin of live fish three, six, and nine months after probiotic exposure. All fish were then euthanized by a Benzocaine overdose followed by a sharp blow to the head, at the nine-month sampling and consequently sampled at three, six, and nine weeks of post-mortem incubation. An inoculation loop was inserted into each bag and then streaked on a blood agar plate containing 0.9 % NaCl. Blood agar plates were incubated at 10 °C for 7 days and subsequently analysed by a skilled technician. Up to two colonies per sample displaying the characteristic yellow Aliivibrio phenotypes (Figure 2) were transferred to fresh blood agar plates to recover monocultures for DNA isolation to determine whether these belonged to the administered species. To ensure consistency and accuracy, colony sampling in trial 1 was performed in a blinded manner by two independent technicians.

**Figure 2:**
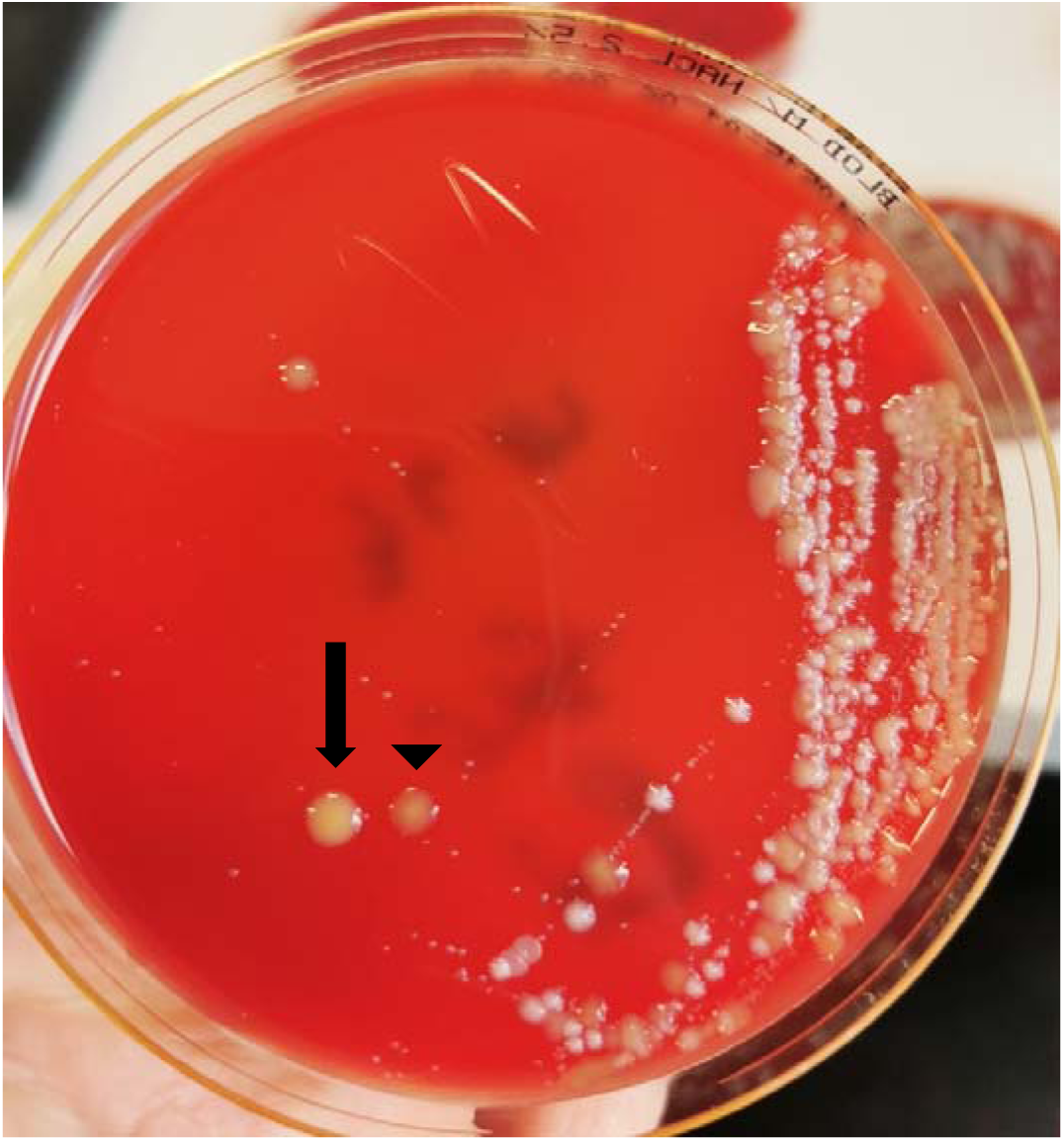
Colonies from the skin of a fish carcass aged for 21 days. Many phenotypic yellow colonies of both administered strains are seen, with single colonies of Aliivibrio sp. strain Vl2 (arrow) and Aliivibrio sp. strain Vl1 (arrowhead) confirmed by qPCR.

In trial 2, sampling was conducted as described for trial 1, but up to 10 colonies per sample were chosen without blinding and subcultured for DNA isolation. The bags in part 2.1 kept at different temperatures were sampled on days 1, 4, and 7 post-mortem while the bags in parts 2.2 and 2.3 were sampled at several timepoints up to day 21 post-mortem (exact timepoints are detailed in the results). In trial 2.3, tissue samples from distal intestines were placed in 10 % buffered formalin at the time of euthanasia and day four post-mortem. At the final sampling in trial 2.3 (21 days post-mortem), samples of skin with underlying muscle from every fish were placed in 10 % buffered formalin and RNA later (Thermo Fisher Scientific, USA). Two fish in trial 2.3 had developed ulcers ≈1 cm in diameter. These ulcers were sampled by swabbing, followed by cultivation at the same timepoints as the other fish. In addition, half of each ulcer was submerged in 10 % phosphate-buffered formalin on day 7.

In a small pilot, we also controlled for the water as a possible source of the administered bacteria as five control fish were dipped in tank water of fish previously exposed to probiotic bacteria, euthanised and incubated for 21 days at 6 °C.

### 2.5 DNA isolation

Following subculture, one inoculation loop with bacterial colony material per isolate was transferred to sterile nuclease-free water (Thermo Fisher Scientific, USA) and incubated at 100 °C for 10 minutes. The samples were then centrifuged at 8000x g for three minutes before the supernatant was diluted 1:5 in water and kept at 4°C until proceeding with qPCR.

Skin and muscle samples kept in RNA-later were weighed and standardized to 25 µg of tissue input for DNA isolation using Blood and tissue kit (Qiagen) following the manufacturer’s instructions. The purity and concentration of DNA were analysed using Multiskan SkyHigh Microplate Spectrophotometer (Thermo Fisher Scientific, USA), and samples with DNA concentrations <10 ng/ml or 230:260 and 260:280 <1.8 were discarded, and DNA isolation was repeated. Finally, all samples were diluted 1:5 before proceeding with qPCR.

### 2.6 qPCR

Both bacterial targets were detected using a Power Track™ SYBR™ green (Thermo Fisher Scientific, Baltics UAB, Lithuania) based qPCR assay with custom-designed oligomers (primers) which target DUF2057 domain-containing protein (615 bp) and hypothetical protein (1542 bp) coding regions of Vl1 and Vl2, respectively. The design of the primers is listed in Table 1.

**Table 1:**
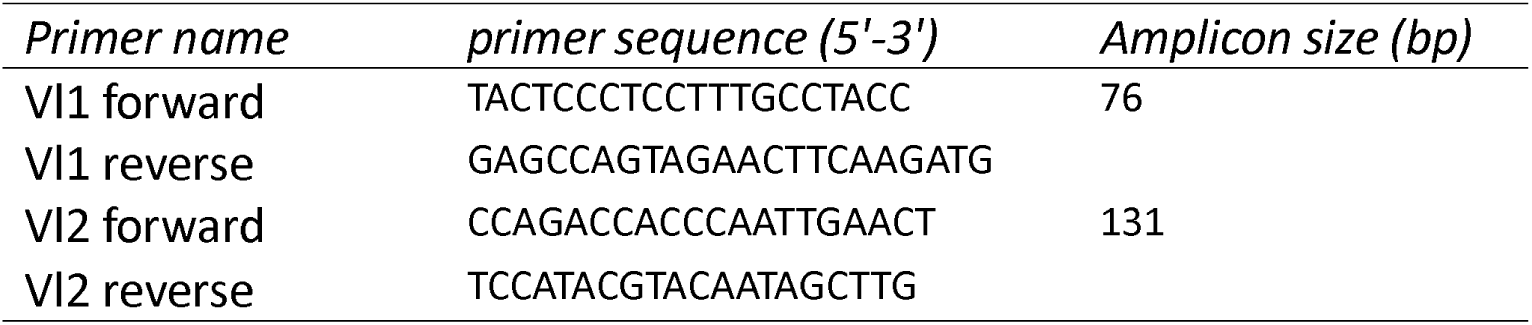
Primer design for qPCR assay targeting the two bacterial strains used in this study.

The qPCR was performed using a total volume of 20 µL for each sample, 10 µL Power Track SYBR green master mix (Thermo Fisher Scientific), 300 nM of each primer, 0.5 µL of 40X Yellow buffer (Thermo Fisher Scientific), 6.5 µL of molecular grade water, and 2 µL of DNA template.

The qPCR assays were carried out in the QuantStudio™ 1 Real-Time PCR System (Applied Biosystems™) using the following parameters: 95^⍰^C for 10 min for initial denaturation followed by 40 cycles of 95^⍰^C for 15 seconds and 60^⍰^C for 60 seconds. Melting curves were generated using thermal conditions of 95^⍰^C for 1 second, 60^⍰^C for 20 seconds, and 95^⍰^C for 1 second, with continuous fluorescence measurements. Positive controls (bacterial DNA template obtained from pure cultures of each strain) and a negative control without DNA were also included in every run. Cycle threshold (Ct) values were recorded, and Ct-values of < 30 were considered positive results. All samples were run in duplicates, discarding and repeating samples with variations of >1 Ct value. Both oligomers demonstrated complete specificity in silico when tested against Atlantic salmon and all known bacterial species by BLAST (Basic local alignment search tool) and qPCR against five different species within the same genus (A. fischeri (At4), A. logei (ATCC15382), A. salmonicida (NCMB2262), Vl1 and Vl2).

### 2.7 Antibodies

Cultures of Vl1 and Vl2 were inactivated in 3 % formalin before the bacterial cells were spun down and added Freunds complete adjuvant. The mixture of antigens and adjuvant (0.5 ml) was injected subcutaneously in a large rabbit as the primary immunization. The first booster dose (0.5 ml) was administered 14 days later with Freunds incomplete adjuvant, followed by the second booster dose 21 days after the primary dose, and a third booster dose 28 days after the primary dose. The second and third booster doses were identical to the first booster dose. Blood samples were collected from the ear vein at each immunization to monitor the development of specific antibodies. Final blood sampling was conducted from the anaesthetized rabbit 42 days after the primary immunization. The collected blood was allowed to clot before centrifugation. Serum containing polyclonal antibodies was aliquoted in 10 ml plastic tubes and stored at -80 °C until use. To decrease cross reactivity before use, the serum was thawed and diluted 1:100 in Da Vinci Green Diluent (Biocare Medical, USA). The serum was then incubated with a strain of Pseudomonas sp. isolated from Atlantic salmon skin for 1 hour, centrifuged at 8000x g for 10 minutes and sterile filtered twice with 0.2 µm pore size Minisart Syringe Filter (Sartorius, Germany). This process was repeated after an additional incubation with the A. wodanis type strain (NCIMB 13582) before use.

### 2.8 Immunohistochemistry

After 24-48 hours, the formalin-fixed skin and muscle samples were dehydrated and subsequently paraffin-embedded. Sections of 5 µm from the formalin-fixed paraffin-embedded (FFPE) samples were placed on glass slides (super-frost plus, Menzel Gläser, Thermo Fisher Scientific) and stored at 4 °C until staining. Glass slides were deparaffinized in xylene and rehydrated through a descending alcohol series. Glass slides were washed in PBS for five minutes, and antigens were unmasked by heat-induced epitope retrieval (HIER) using a pressure cooker at 110 °C with DIVA Decloaker (Biocare Medical) for 10 minutes. The staining was carried out using a MACH 1 universal HRP polymer detection kit (Biocare Medical) following the manufacturer’s instructions. The tissue samples were incubated with the primary antibodies in 1:10 000 dilution for one hour. Optimal antibody concentration for a positive signal with an absence of cross-reaction was determined using un-incubated and incubated controls from trial 2 and positive control samples containing Vibrio anguillarum and Aliivibrio salmonicida supplied by the Norwegian Veterinary Institute.

### 2.9 Modified Gram-staining

FFPE ulcer samples were prepared as described for immunohistochemistry but instead stained with modified Gram-staining, using Hucker-Conn, acetone, Lugol’s iodine, and neutral red for one minute.

### 2.10 16S *rDNA* gene sequencing

To control for the natural exposure to unadministered Aliivibrio spp., twelve isolates were selected for typing using 16S rDNA gene sequencing. These isolates were chosen based on the phenotypical similarity (Figure 2) to the administered strains while being qPCR negative when tested for the Vl1 and Vl2 oligomers. Ten samples of DNA isolated from representative colonies in trial 1 and two samples of DNA isolated from representable colonies in trial 2.2 were submitted to Eurofins Genomics (Germany) for 16 S rRNA gene Sanger sequencing.

### 2.11 Statistical analyses

Sample sizes were chosen based on the number of available fish and previous pilot studies. Chi-square (X^2^) was used to compare skin and muscle samples (positive and negative for the administered strains) by qPCR in different groups in trial 2.3 at 21 days post-mortem (n=16). All assumptions for this statistical method were met.

## 3. Results

This study aimed to investigate whether these bacteria could colonize Atlantic salmon after administration, determine the tissue tropism and assess the duration of colonization. An overview of all results produced in this study is presented in Table 2.

**Table 2:**
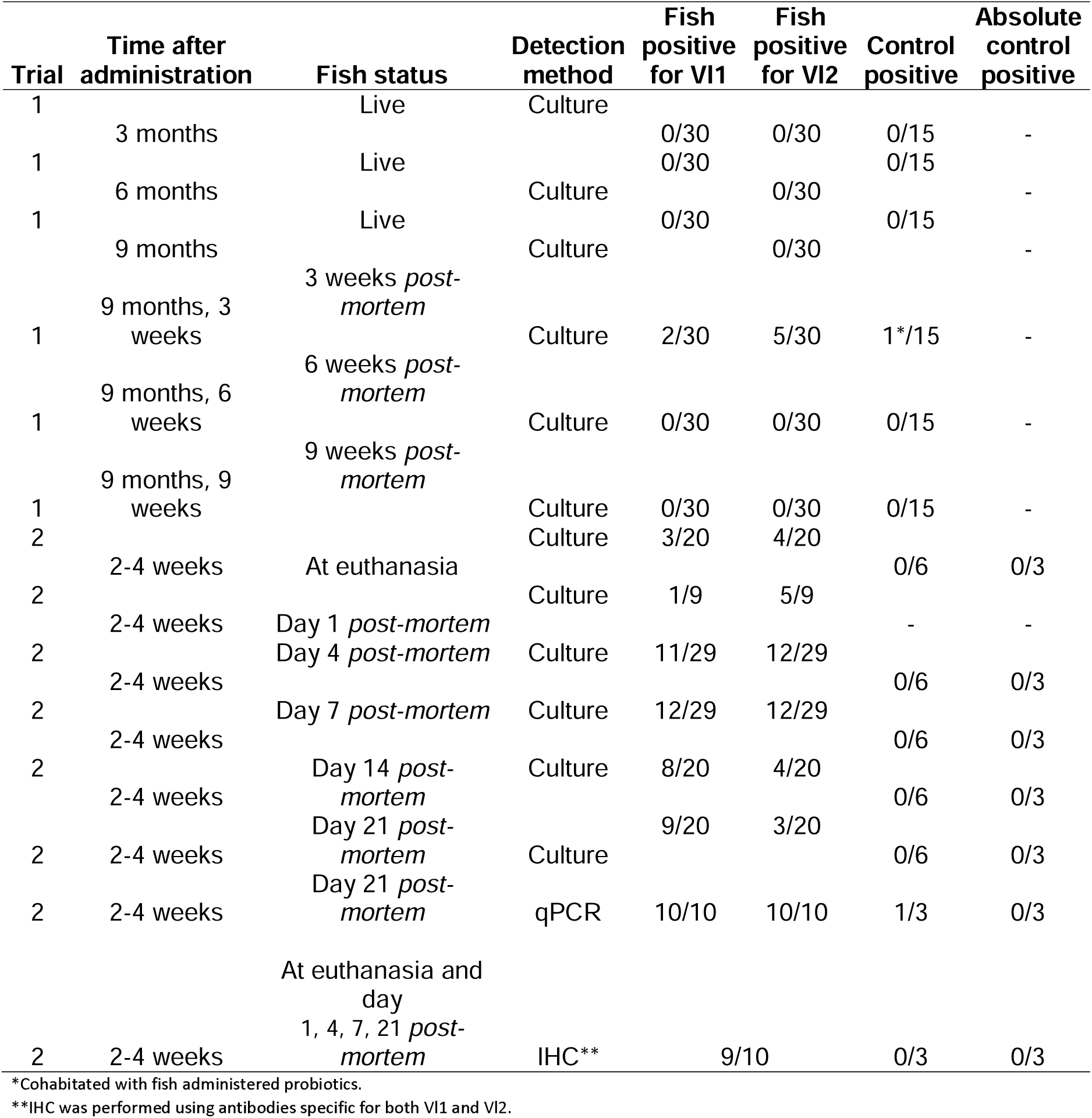
Overview of every sampling with all fish positive for the administered probiotics (N=83).

### 3.1 Culture

In trial 1, we aimed to re-isolate the administered strains from the skin of live fish over a 9-month period following probiotic administration. However, the strains were not detected on any sampled timepoint. In contrast, incubation of whole fish post-mortem (after euthanasia 9 months post-administration) allowed for the detection of the probiotic strains. Probiotic Aliivibrio colonies were only retrieved at 3 weeks post-mortem. At this timepoint, a total of 55 colonies were sampled with 8 (14.5 %) being identified, as one of the two administered strains by qPCR. In the probiotic-administered group, Vl1 was detected in two fish (6.7 %) and Vl2 was detected in five fish (16.7 %). Additionally, Aliivibrio sp. strain Vl2 was detected in one cohabitant fish (10 %) at the same timepoint. Neither probiotic strain was detected in the control fish kept in a separate tank (0 %). While the majority of sampled colonies tested negative by qPCR, 16S rDNA sequencing confirmed that all ten selected qPCR negative colonies were other Aliivibrio spp., as identified by BLAST search and an alignment against a custom 16S rRNA gene library containing sequences of Vl1 and Vl2 (Table S6).

As we were able to re-isolate the administered strains in trial 1, the next aim was to investigate the effect of temperature on re-isolation and tissue location of the probiotics. In trial 2.1, nine fish administered probiotics were euthanised two weeks after probiotic immersion and sampled on days 1, 4 and 7 post-mortem. Across all timepoints, the administered species were re-isolated from all samples incubated at 6°C and 10°C, but none at 4°C (Figure 3, Table S1). The re-isolation from 6°C and 10°C were almost identical, with the exception of one more positive fish at 6°C, day 1 post-mortem. A total of 141 colonies were sampled, and 100 % were qPCR-positive for the administered strains (27.7 % Vl1 and 72.3 % Vl2). Overall, Vl1 was recovered from 11.1 %, 55.6 % and 33.3 % of fish at day 1, 4 and 7 days post-mortem, respectively. Vl2 was recovered from 55.6 % of fish at every timepoint post-mortem. Colonies identified as the administered strains were recovered from both skin and the pooled visceral organs.

**Figure 3:**
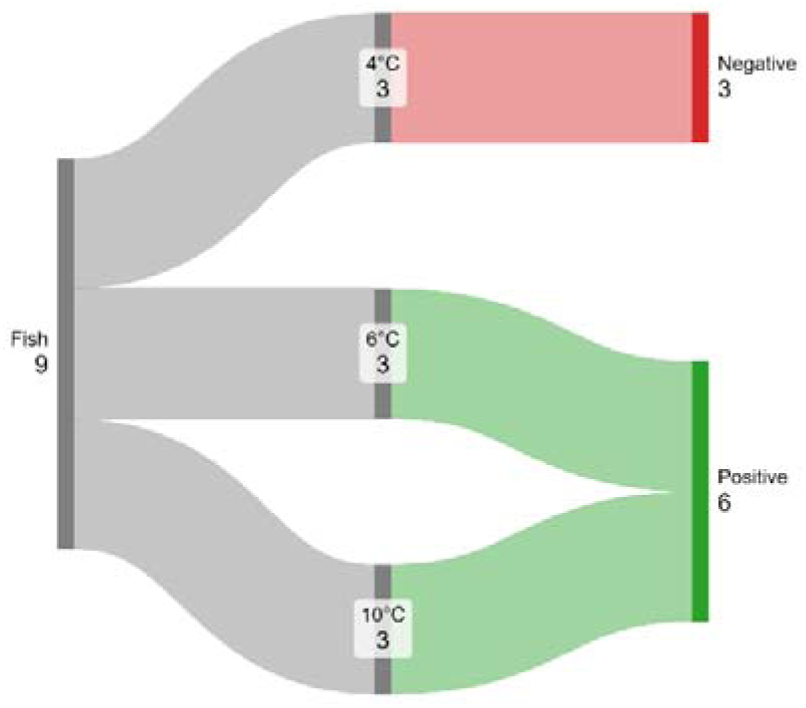
Sankey plot showing re-isolation of administered Aliivibrio species at different temperatures in trial 2.1, identified by qPCR (n=9). Figure made with sankeymatic.com.

Next, we aimed to investigate tissue location using a larger number of fish at 6°C for probiotic recovery. In trial 2.2, 10 fish administered probiotics and 3 control fish were euthanised three weeks after probiotic or mock immersion and sampled immediately (at the start of incubation) and on days 1, 2, 3, 4, 7, 14, and 21 post-mortem. A total of 111 colonies were retrieved for identity confirmation by qPCR; however, only one colony was positive for the administered strains. This colony was isolated on day 7 post-mortem from the visceral organs of one fish that had received probiotics (Table S2). Additionally, two qPCR-negative colonies were identified as Aliivibrio sp. through 16S rDNA sequencing and BLAST analysis (Table S6).

Finally, we separated the carcass from the distal intestine and remaining visceral organs to investigate which specific tissues were colonized by the probiotics. In trial 2.3, 10 fish administered probiotics and 3 control fish were euthanised four weeks after probiotic or mock immersion and sampled immediately (at the start of incubation) as well as on days 4, 7, 14, and 21 post-mortem. Viable Aliivibrio spp. were recovered from the skin, visceral organs, and distal intestine at all timepoints, except for the distal intestine on day 21 (Figure 4, Table S3). A total of 257 colonies were sampled, of which 160 (62,3 %) were identified by qPCR as one of the two administered strains. None of the colonies isolated from the control fish were identified as either of the two administered strains. The percentage of fish from which the probiotics were successfully reisolated peaked on day 7 post-mortem across all tissue samples. Re-isolated Aliivibrio colonies from the visceral organs and distal intestine decreased after day 7, whereas positive cultures from skin remained high throughout the experiment. The administered probiotics were most frequently re-isolated from the skin at all timepoints, with the exception of day 4 post-mortem.

**Figure 4:**
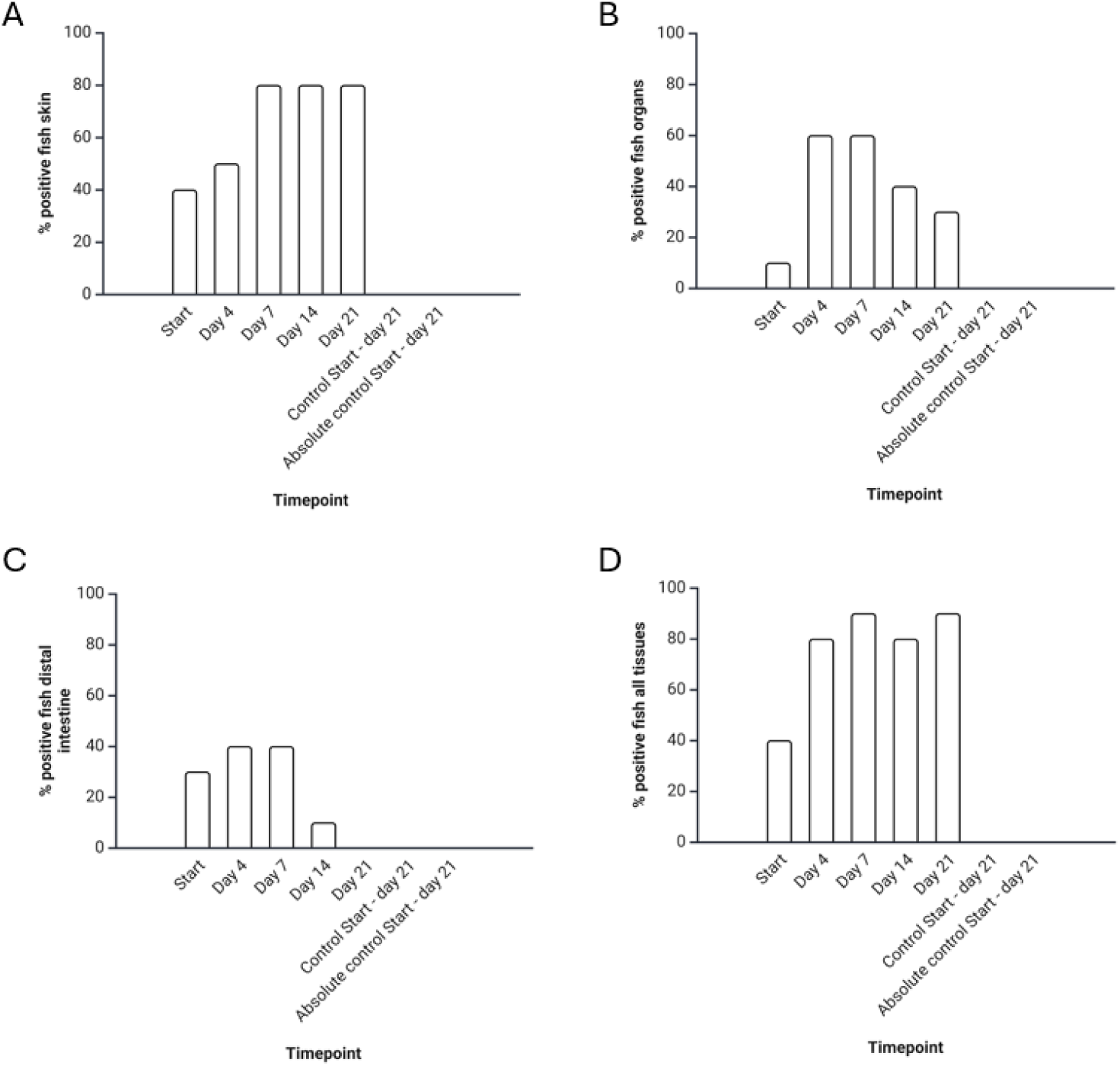
Percentage of fish sampled at different timepoints after euthanasia in trial 2.3 where administered probiotic bacterial colonies were recovered (10 immersed, 3 controls, 3 absolute controls, n=16). Absolute controls were euthanized before arrival at the research facility to avoid contact with saltwater, while controls were mock immersed and euthanized at the same time as fish administered probiotics. Three locations were swabbed and cultivated: A) Skin, B) visceral organs without distal intestine and C) distal intestine. All tissues combined are also presented D). Figure made with Biorender.com.

After sampling from fish tissues and organs, particularly following incubation of the carcasses, a diverse range of bacteria grew on the agar plates. The types and numbers of bacterial species cultivated varied depending on the sampling time. At its peak in trial 2, colonies phenotypically similar to the administered Aliivibrio spp. only amounted to roughly 20 % of the total cultured colonies. In contrast, nearly 100 % of the cultured colonies in trial 1 resembled the administered Aliivibrio spp. at three weeks of post-mortem incubation. Also, there was a notable difference between the recovery rate as only 14.5 % of the colonies sampled in trial 1 were of the administered strains compared to 100 %, 0.9 % and 62.3 % in trial 2.1, 2.2 and 2.3, respectively. All five control fish in the small pilot remained negative for the administered strains during 21 days of post-mortem incubation.

### 3.2 qPCR analyses of skin and muscle samples at day 21 *post-mortem*

Next, we sought to quantify bacterial colonization in fish administered probiotics versus controls using qPCR on tissues of skin and muscle. All 10 fish from trial 2.3 that had been administered probiotics and were incubated for 21 days post-mortem tested positive for the administered strains by qPCR (Figure 5, Table S4). One of three control fish and none of the absolute controls were positive for the administered strains by qPCR at 21 days post-mortem. A chi-square test was performed to examine the difference in qPCR positive fish in the group administered probiotics versus the controls. The difference between these groups was significant, X (1, N = 13) = 7.88, p = .005, fish administered probiotic Aliivibrios four weeks earlier had higher levels of the probiotic bacteria in the skin and muscle samples after 21 days of post-mortem incubation.

**Figure 5:**
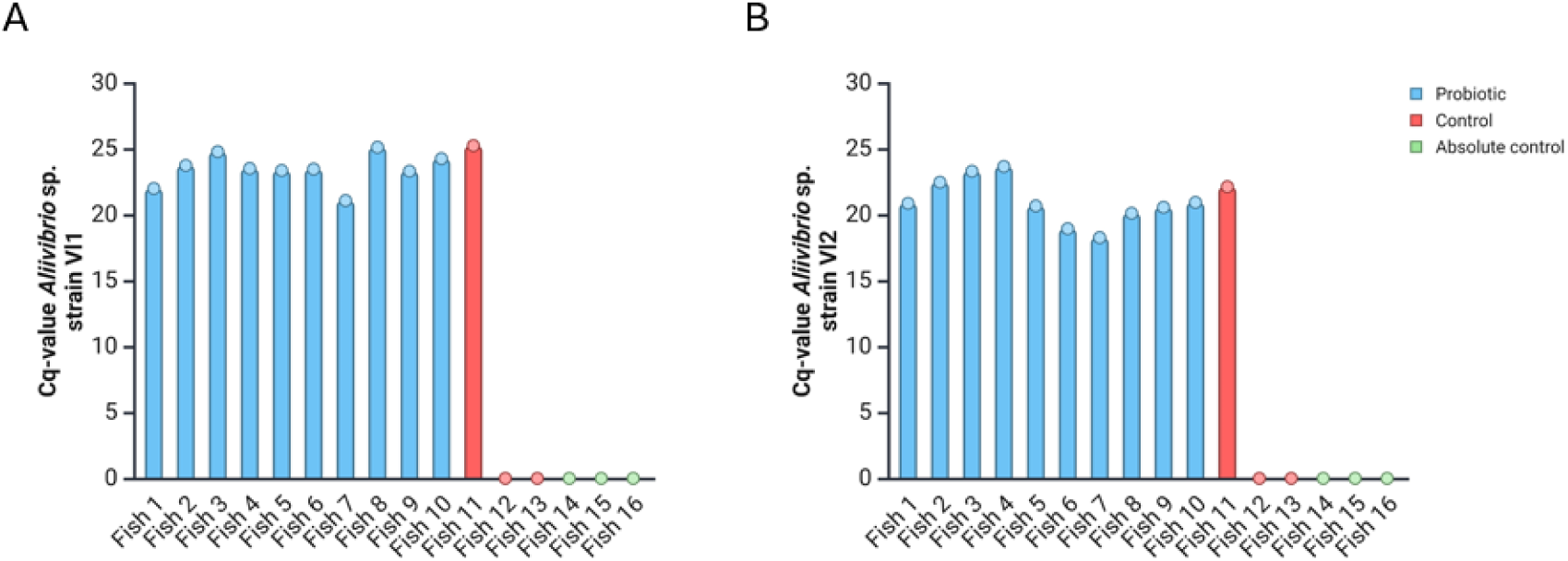
qPCR results for samples of skin with muscle (n=16) at 21 days post euthanasia, using Vl1 (A) and Vl2 (B) primers.

### 3.3 Immunohistochemistry

To visualize the location of probiotic bacteria, immunohistochemistry staining was performed on fish tissues. At day 21 post-mortem, all fish previously immersed in probiotics exhibited positive reactions, with structures resembling bacterial cells appearing as red stains in the skin and muscle tissues. In contrast, no similar reactions were observed in controls or absolute controls. Positive cells were detected throughout the upper layers of the epidermis, as well as in the dermis, hypodermis and muscle tissue (Figure 6).

**Figure 6:**
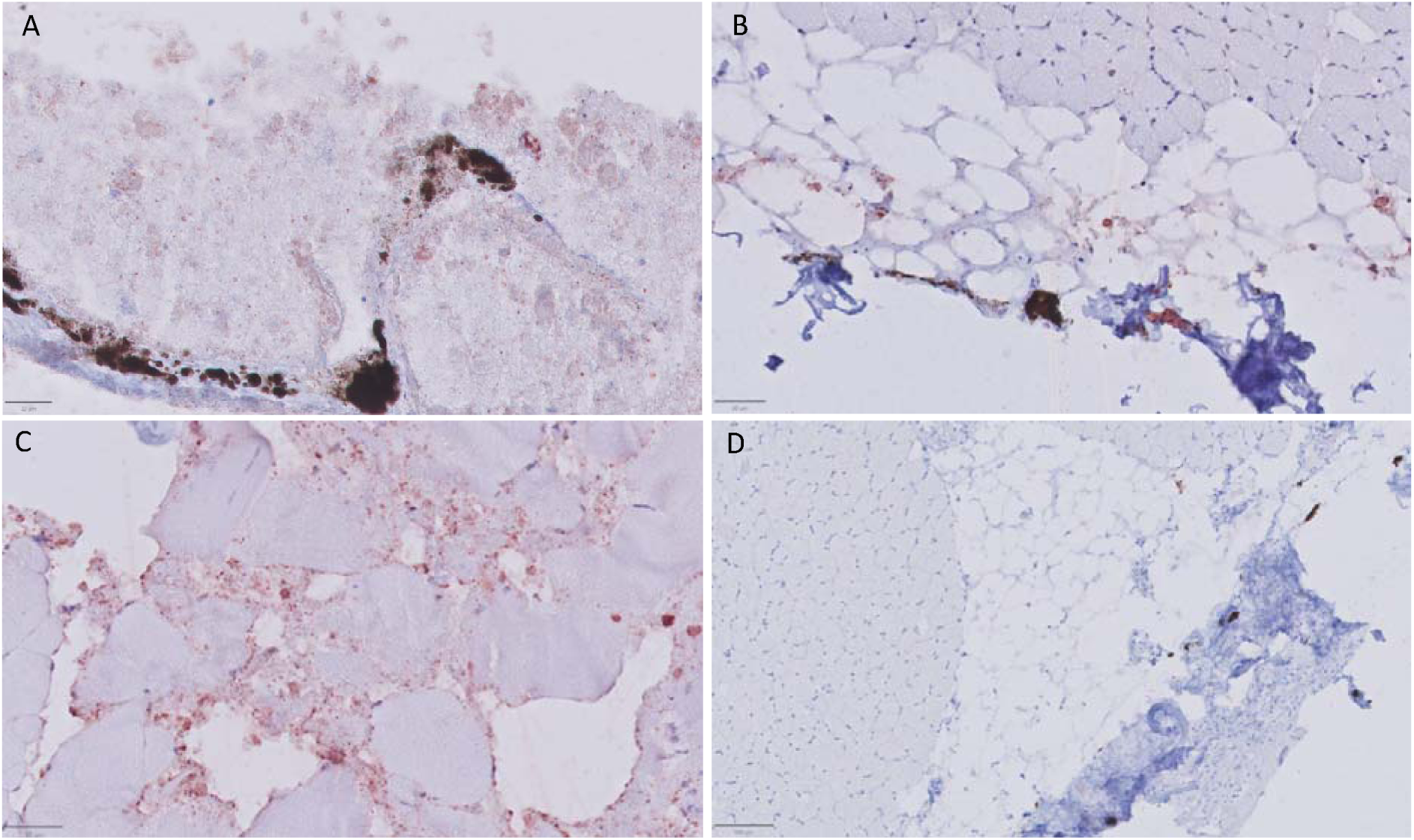
Immunohistochemistry in representative layers of skin and muscle after 21 days of aging in trial 2.3 (scale bars = 20-100 µm). A) epidermis, B) dermis and hypodermis, C) muscle, and D) control sample including epidermis, dermis, hypodermis and muscle tissue. Red stains indicate the presence of the administered Aliivibrio species.

There were also positive cells visualized in samples taken from the distal intestine at the time of euthanasia and day four of aging. The control and absolute control samples were negative by IHC at all timepoints (Figure 7).

**Figure 7:**
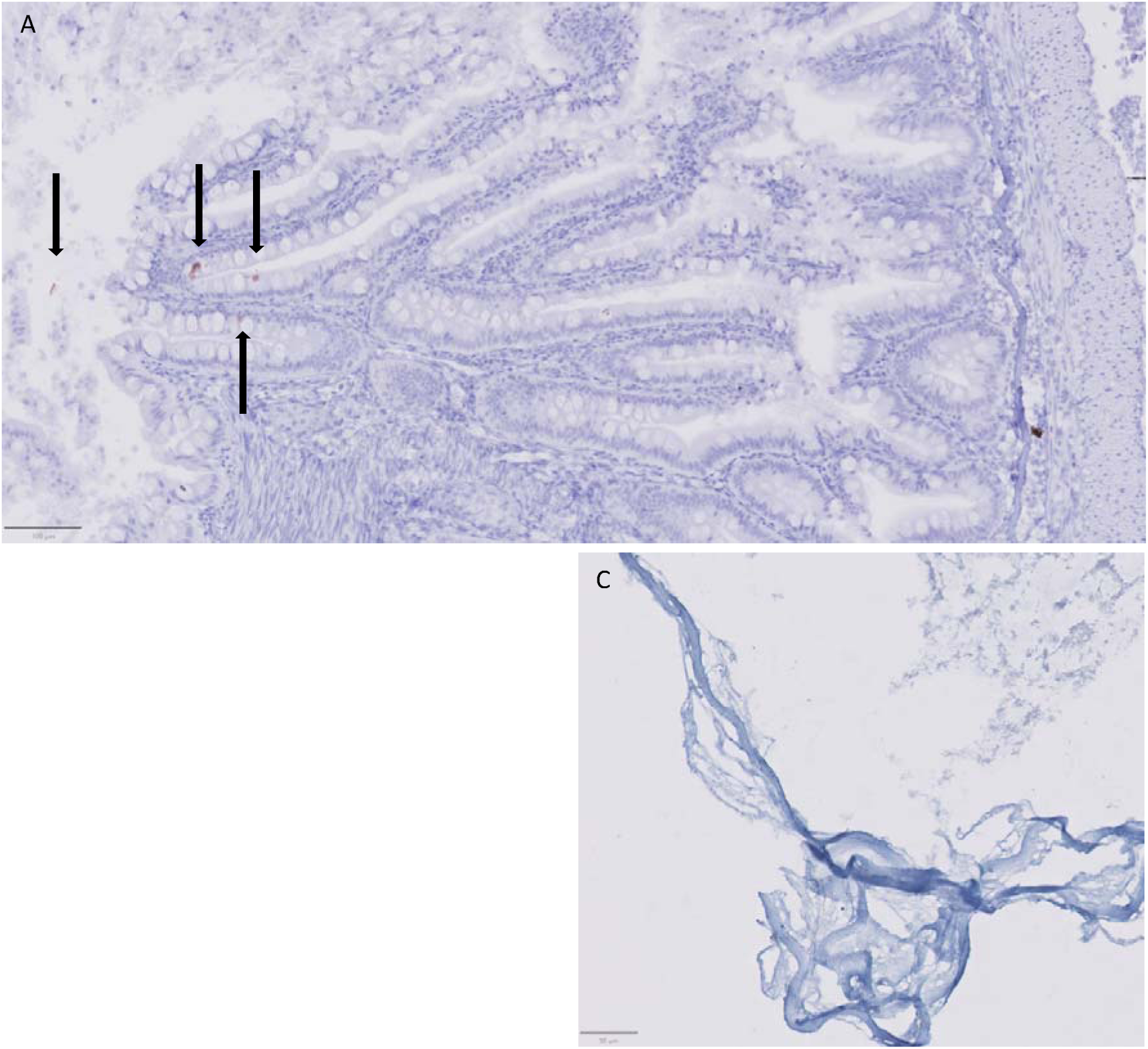
Immunohistochemistry of representative samples in the distal intestine in trial 2.3 (scale bars = 50-100 µm). A) Four positive colonies (arrows) visualized in a sample taken at the time of euthanasia, B) a loop of stratum compactum (sc) harbouring a larger positive colony at 4 days of incubation and C) negative control.

### 3.4 Ulcers

A mixed bacterial community including the administered Aliivibrio spp. was recovered from both ulcers in trial 2.3. Although colonies identified as the probiotics were found in both ulcers at the start, the number of colonies resembling these species only amounted to < 10 % of the total bacteria at this time. This number increased during aging to ≈30 % (±10 %) at day 21. Colonies identified as the administered Aliivibrio spp. were reisolated from at least one of two ulcers at every timepoint except for day 4 and visualized by IHC and modified Gram-staining (Figure 8, Table S5).

**Figure 8:**
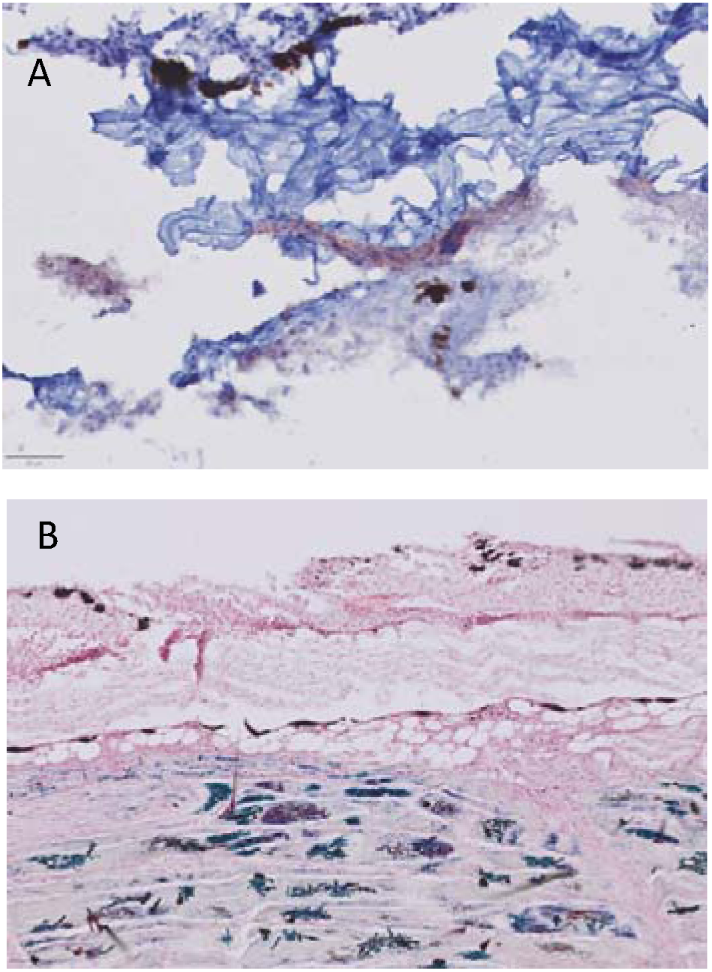
A) Ulcers from trial 2.3 with positive cells by immunohistochemistry in the dermis layer at seven days of ageing, B) modified Gram-staining of the same sample shows pockets of larger quantities of Gram-positive and -negative bacteria in all layers of the skin, primarily in the muscle-layer. Scale bars represent actual size.

## 4. Discussion

In this study, we demonstrated the successful colonization of Atlantic salmon by two probiotic Aliivibrio strains up to nine months after bath administration. Both strains were re-isolated through culture and detected using qPCR and immunohistochemistry. While probiotic bacteria were detectable in fish during the first four weeks post-administration, a period of post-mortem incubation significantly increased the detection rate. Most of the decaying fish rendered growth of the probiotic bacteria from the skin by day 21, with the highest recovery rates for the distal intestine and remaining visceral organs observed on day 4 and 7. Considering all sampled organs combined, day 7 and 21 were the most effective sampling points, with the administered bacteria recovered post-mortem from 90 % of the fish immersed in probiotics, compared to only 40 % at the start of the incubation period. The pilot controlling for presence of probiotics in the rearing water indicates that the recovered bacteria in these trials are from the fish and not bacterial residues from the probiotic administration residing in the tank water. This is in-line with other studies with similar design^74^.

The increased recovery of these Aliivibrio species during post-mortem incubation of fish carcasses may result from the growth of either (1) viable bacteria missed during initial sampling(s), or (2) VBNC bacteria transitioning to culturable states. In trial 1, no viable administered Aliivibrio bacteria were detected in any fish before post-mortem incubation. This suggests that VBNC bacteria may have re-entered culturable states when the nutrients of the carcass became available, as the need for dormant survival mechanisms were no longer necessary. This phenomenon is well-documented in other Vibrio species ^50,53^. However, the administered Aliivibrio strains were re-isolated from several fish before post-mortem incubation in trial 2.3, indicating that the probiotics can also persist in culturable states within the live host. Regardless of the mechanism, the aging carcass appeared to serve as a favourable enrichment medium, as the colonized strains exhibited robust growth during post-mortem incubation in both trials.

The effects of bath-applied probiotic Aliivibrio spp. have been shown to benefit skin health in Atlantic salmon and lumpfish, demonstrated by a reduction in ulcer prevalence ^75,76^. One commonly described mechanism for probiotic bacteria is colonization resistance via competitive exclusion ^78^. The ability of the administered strains to colonize both the skin and ulcers, as shown in our results, leaves competitive exclusion as a possible mechanisms of action for these probiotic strains. A. wodanis is closely related to the probiotics administered in this trial and this species is documented to strongly out-compete the primary ulcer pathogen M. viscosa through bacteriocin and siderophore production ^79^. While this results in reduced acute mortality, the disease transitions to a more chronic ulcerative condition with prolonged pathogenesis ^34^. In contrast, the probiotic strains used here are non-pathogenic, as demonstrated by Klakegg et al. who observed reduced mortality following probiotic bath administration ^75^. This suggests that the probiotic strains do not prolong the disease, although the competitive exclusion mechanisms may be similar or genus specific.

Probiotic strains were also detected in the distal intestine, with up to 30 % of fish harbouring the bacteria at the time of euthanasia. Since rearing water can enter the gut of fish passively, large quantities of viable bacteria in the water likely facilitate their entry and colonization of the fish intestinal tract. Colonies representing the administered Aliivibrio strains were also found within the remaining visceral organs, which can be explained by the middle intestine being part of the visceral organs, as Aliivibrio spp. has previously been shown to reside here ^40^. Still, other tissues in the remaining visceral organs cannot be ruled out as possible sources. The presence of viable Aliivibrio species weeks or months after administration from the intestine and skin, suggests the possibility of a self-enforcing biological ecosystem; probiotic bacteria shed from the skin could enter the intestinal tract of the fish through water, while bacterial cells from the gut expelled with faeces can settle on the skin. The detection of the administered strains on the skin of a cohabitant control in trial 1 supports this hypothesis. This has also previously been suggested by Karlsen et al., who found an increase in the number of Aliivibrio cells in the intestine of fish with ulcers ^39^. Re-isolation from the intestine aligns with the findings of Klakegg et al. linking probiotic Aliivibrio spp. to improved feed conversion rates (FCR), suggesting that intestinal colonization by these bacteria may directly influence the digestive system, consistent with studies on numerous probiotics and fish species ^26^.

Trial 2 highlights the dynamic nature of the microbiome, with 100 % of the sampled colonies testing positive by qPCR at 2 weeks post-administration. The number dropped sharpy to 0.9 % at 3 weeks post-administration before rising again to 62.3 % at 4 weeks post-administration. All qPCR-negative colonies from trials 1 and 2 that were analysed, were confirmed by 16S rDNA gene sequencing to be other Aliivibrio species. This suggests that the low recovery rate observed 3 weeks after administration was due to an influx of transient non-administered species, naturally acquired. Furthermore, it accounts for why only 7.8% of sampled colonies in trial 1 tested positive by qPCR, as nine months in seawater likely introduced the fish to more phenotypically similar Aliivibrio bacteria.

Natural exposure to various Aliivibrio spp. in seawater was also evident in one control fish from trial 2.3, which tested qPCR-positive for both probiotic strains despite not being administered the probiotics. While we cannot exclude the possibility of false positives from unknown field strains with similar genes, this seems unlikely as the remaining five control fish were negative by qPCR, and all control fish yielded negative results by IHC and re-isolation. This demonstrates that the sensitivity of qPCR in this trial was superior compared to culturing and IHC, which is one of the primary strengths of this technology ^80–85^. The added advantage of detection by IHC is the ability to visualize the bacteria in situ. Positive cells were identified in all skin-muscle tissue layers, and the bacteria appeared to be most abundant in the muscle tissue. Interestingly, the bacteria seem to follow the perimysium from the dermis into the muscle tissue and reside in the intermuscular tissue. The Aliivibrio bacteria do not appear on top of any muscle fibres, but almost exclusively between them.

In conclusion, we successfully re-isolated viable probiotic bacteria up to nine months after a single bath administration, showing a long-lasting colonization profile. The findings presented in this study are of great interest because the health benefits conferred by probiotic bacteria are often linked to their ability to colonize and persist within the host ^12,86–91^. The bacteria were capable of colonizing the skin, visceral organs, distal intestine, and ulcers, and we detected the administered strains in fresh and post-mortem incubated fish by culture. We also applied methods for detection of both Aliivibrio strains using qPCR and for visualisation of both strains in situ by immunohistochemistry. The use of three complementary methods confirms that both Aliivibrio spp. can colonize Atlantic salmon following bath administration. The detection of administered strains in ulcers, combined with previous evidence of ulcer reduction from these probiotics, supports competitive exclusion as a potential mode of action. Further studies are warranted to investigate how these probiotic bacteria influence ulcerative conditions in Atlantic salmon and lumpfish.

## Supplementary materials

All data is available in the supplementary materials: Table S1, Re-isolation of administered Aliivibrio species at different temperatures in trial 2.1, confirmed by qPCR (n=9); Table S2, Re-isolation of administered Aliivibrio species in trial 2.2, confirmed by qPCR (n=13), Table S3, Re-isolation of administered Aliivibrio species in trial 2.3, confirmed by qPCR (n=16).; Table S4, Ct-values for both administered strains in DNA-samples isolated from skin per fish and group average at 21 days post euthanasia (n=16); Table S5, Overview of ulcers positive for both administered strains recovered by culturing at different timepoints in trial 2.3 (n=2); Table S6, 16S rDNA sequences from qPCR negative colonies resembling the administered strains.

## Author contributions

Marius Steen Dobloug: Conceptualization, Data curation, Formal analysis, Investigation, Methodology, Visualization, Writing – original draft, Writing – review and editing. Stanislav Iakhno: Investigation, Methodology, Writing – review and editing. Simen Foyn Nørstebø: Methodology, Supervision, writing – review and editing. Øystein Evensen: Methodology, Supervision, writing – review and editing. Henning Sørum: Conceptualization, Investigation, Methodology, Supervision, Writing – review and editing.

## Acknowledgements

Thanks to Aud Kari Fauske for preparation of the antibodies used in this trial and Camilla Skagen-Sandvik for practical help with the fish during both trials. Thanks to Stein Helge Skjelde (Sørsmolt) and Fister Smolt for the fish used in this trial.

