## Supplementary materials for "Long-Term Colonization Dynamics of Probiotic *Aliivibrio* spp. in Atlantic Salmon (*Salmo salar) Following Bath Administration*"

Table S1: Re-isolation of administered *Aliivibrio* species at different temperatures in trial 2.1, confirmed by qPCR (n=9).

| **Tissue** | **Temperature** | **Timepoint** | **% Positive A. njordis** | **% Positive A. balderis** | **% Positive A. njordis or A. balderis** |
| --- | --- | --- | --- | --- | --- |
| Skin (Fish 1-3) | 4 °C | Day 1 | 0 % | 0 % | 0 % |
| Skin (Fish 1-3) | 4 °C | Day 4 | 0 % | 0 % | 0 % |
| Skin (Fish 1-3) | 4 °C | Day 7 | 0 % | 0 % | 0 % |
| Skin (Fish 4-6) | 6 °C | Day 1 | 33 % | 100 % | 100 % |
| Skin (Fish 4-6) | 6 °C | Day 4 | 100 % | 66 % | 100 % |
| Skin (Fish 4-6) | 6 °C | Day 7 | 66 % | 66 % | 100 % |
| Skin (Fish 7-9) | 10 °C | Day 1 | 0 % | 66 % | 66 % |
| Skin (Fish 7-9) | 10 °C | Day 4 | 66 % | 100 % | 100 % |
| Skin (Fish 7-9) | 10 °C | Day 7 | 66 % | 100 % | 100 % |
| Abdominal organs (Fish 1-3) | 4 °C | Day 1 | 0 % | 0 % | 0 % |
| Abdominal organs (Fish 1-3) | 4 °C | Day 4 | 0 % | 0 % | 0 % |
| Abdominal organs (Fish 1-3) | 4 °C | Day 7 | 0 % | 0 % | 0 % |
| Abdominal organs (Fish 4-6) | 6 °C | Day 1 | 0 % | 0 % | 0 % |
| Abdominal organs (Fish 4-6) | 6 °C | Day 4 | 100 % | 100 % | 100 % |
| Abdominal organs (Fish 4-6) | 6 °C | Day 7 | 0 % | 100 % | 100 % |
| Abdominal organs (Fish 7-9) | 10 °C | Day 1 | 0 % | 0 % | 0 % |
| Abdominal organs (Fish 7-9) | 10 °C | Day 4 | 0 % | 100 % | 100 % |
| Abdominal organs (Fish 7-9) | 10 °C | Day 7 | 100 % | 100 % | 100 % |

Table S2: Re-isolation of administered *Aliivibrio* species in trial 2.2, confirmed by qPCR (n=13).

| **Tissue** | **Timepoint** | **% Positive A. njordis** | **% Positive A. balderis** | **% Positive A. njordis or A. balderis** |
| --- | --- | --- | --- | --- |
| Skin | Start | 0 % | 0 % | 0 % |
| Skin | Day 1 | 0 % | 0 % | 0 % |
| Skin | Day 2 | 0 % | 0 % | 0 % |
| Skin | Day 3 | 0 % | 0 % | 0 % |
| Skin | Day 4 | 0 % | 0 % | 0 % |
| Skin | Day 7 | 0 % | 0 % | 0 % |
| Skin | Day 14 | 0 % | 0 % | 0 % |
| Skin | Day 21 | 0 % | 0 % | 0 % |
| Control skin 1-3 | Start-day 21 | 0 % | 0 % | 0 % |
| Abdominal organs | Start | 0 % | 0 % | 0 % |
| Abdominal organs | Day 1 | 0 % | 0 % | 0 % |
| Abdominal organs | Day 2 | 0 % | 0 % | 0 % |
| Abdominal organs | Day 3 | 0 % | 0 % | 0 % |
| Abdominal organs | Day 4 | 0 % | 0 % | 0 % |
| Abdominal organs | Day 7 | 0 % | 6 % | 6 % |
| Abdominal organs | Day 14 | 0 % | 0 % | 0 % |
| Abdominal organs | Day 21 | 0 % | 0 % | 0 % |
| Control abdominal organs 1-3 | Start-day 21 | 0 % | 0 % | 0 % |

Table S3: Re-isolation of administered *Aliivibrio* species in trial 2.3, confirmed by qPCR (n=16).

| **Tissue** | **Time after euthanasia** | **% Positive A. njordis** | **% Positive A. balderis** | **% Positive A. njordis or A. balderis** |
| --- | --- | --- | --- | --- |
| Skin | Start | 20 % | 30 % | 40 % |
| Skin | Day 4 | 40 % | 20 % | 50 % |
| Skin | Day 7 | 80 % | 30 % | 80 % |
| Skin | Day 14 | 80 % | 20 % | 80 % |
| Skin | Day 21 | 80 % | 20 % | 80 % |
| Skin control | Start-day 21 | 0 % | 0 % | 0 % |
| Abdominal organs (without distal intestine) | Start | 10 % | 10 % | 10 % |
| Abdominal organs (without distal intestine) | Day 4 | 40 % | 50 % | 60 % |
| Abdominal organs (without distal intestine) | Day 7 | 50 % | 40 % | 60 % |
| Abdominal organs (without distal intestine) | Day 14 | 30 % | 30 % | 40 % |
| Abdominal organs (without distal intestine) | Day 21 | 30 % | 10 % | 30 % |
| Abdominal organs (without distal intestine) control | Start-day 21 | 0 % | 0 % | 0 % |
| Distal intestine | Start | 30 % | 20 % | 30 % |
| Distal intestine | Day 4 | 20 % | 30 % | 40 % |
| Distal intestine | Day 7 | 30 % | 30 % | 40 % |
| Distal intestine | Day 14 | 10 % | 0 % | 10 % |
| Distal intestine | Day 21 | 0 % | 0 % | 0 % |
| Distal intestine control | Start-day 21 | 0 % | 0 % | 0 % |
| Absolute control (whole fish). | Start-day 21 | 0 % | 0 % | 0 % |

Table S4: Ct-values for both administered strains in DNA-samples isolated from skin per fish and group average at 21 days post euthanasia (n=16).

| **Fish** | **Ct-value A. njordis** | **Ct-value A. balderis** | **Average Ct-value A. njordis** | **Average Ct-value A. balderis** |
| --- | --- | --- | --- | --- |
| 1 | 21,99 | 20,865 |  |  |
| 2 | 23,74 | 22,465 |  |  |
| 3 | 24,78 | 23,3 |  |  |
| 4 | 23,51 | 23,66 |  |  |
| 5 | 23,36 | 20,67 |  |  |
| 6 | 23,46 | 18,94 |  |  |
| 7 | 21,075 | 18,265 |  |  |
| 8 | 25,115 | 20,115 |  |  |
| 9 | 23,305 | 20,565 |  |  |
| 10 | 24,255 | 20,935 | **23,459** | **20,978** |
| Control 1 | 25,24 | 22,125 |  |  |
| Control 2 | Negative | Negative |  |  |
| Control 3 | Negative | Negative | **25,24*** | **22,125*** |
| Absolute Control 1 | Negative | Negative |  |  |
| Absolute Control 2 | Negative | Negative |  |  |
| Absolute Control 3 | Negative | Negative | **Negative** | **Negative** |

*Two out of three samples were negative and therefore not included in the average.

Table S5: Overview of ulcers positive for both administered strains recovered by culturing at different timepoints in trial 2.3 (n=2).

| **Location** | **Timepoint after euthanasia** | **% Positive A. njordis** | **% Positive A. balderis** | **% Positive A. njordis or A. balderis** |
| --- | --- | --- | --- | --- |
| Ulcer | Start | 100 % | 50 % | 100 % |
| Ulcer | Day 4 | 0 % | 0 % | 0 % |
| Ulcer | Day 7 | 50 % | 50 % | 50 % |
| Ulcer | Day 14 | 50 % | 0 % | 50 % |
| Ulcer | Day 21 | 100 % | 0 % | 100 % |

Table S6: 16S rDNA sequences from qPCR negative colonies resembling the administered strains.

| **Trial 1, colony 1** | **Sequence** | |
| --- | --- | --- |
| Trial 1, colony 2 | | GTAGCTTGCTACTTTGCTGACGAGCGGCGGACGGGTGAGTAATGCCTGGGAATATGCCTTGATGTGGGGGATAACTATTGGAAACGATAGCTAATACCGCATAATGTCTTCGGACCAAAGAGGGGGATCTTCGGACCTCTCGCGTCAAGATTAGCCCAGGTGAGATTAGCTAGTTGGTGGGGTAAGAGCTCACCAAGGCGACGATCTCTAGCTGGTCTGAGAGGATGATCAGCCACACTGGAACTGAGACACGGTCCAGACTCCTACGGGAGGCAGCAGTGGGGAATATTGCACAATGGGCGAAAGCCTGATGCAGCCATGCCGCGTGTATGAAGAAGGCCTTCGGGTTGTAAAGTACTTTCAGTCGTGAGGAAGGGTGTGCAGTTAATAGCTGCATATCTTGACGTTAGCGACAGAAGAAGCACCGGCTAACTCCGTGCCAGCAGCCGCGGTAATACGGAGGGTGCGAGCGTTAATCGGAATTACTGGGCGTAAAGCGCATGCAGGTGGTTCATTAAGTCAGATGTGAAAGCCCGGGGCTCAACCTCGGAACCGCATTTGAAACTGGTGAACTAGAGTGCTGTAGAGGGGGGTAGAATTTCAGGTGTAGCGGTGAAATGCGTAGAGATCTGAAGGAATACCAGTGGCGAAGGCGGCCCCCTGGACAGACACTGACACTCAGATGCGAAAGCGTGGGGAGCAAACAGGATTAGATACCCTGGTAGTCCACGCCGTAAACGATGTCTACTTGGAGGTTGTGGCCTTGAGCCGTGGCTTTCGGAGCTAACGCGTTAAGTAGACCGCCTGGGGAGTACGGTCGCAAGATTAAAACTCAAATGAATTGACGGGGGCCCGCACAAGCGGTGGAGCATGTGGTTTAATTCGATGCAACGCGAAGAACCTTACCTACTCTTGACATCTACAGAATTCGCTAGAGATAGCTTAGTGCCTTCGGGAACTGTAAGACAGGTGCTGCATGGCTGTCGTCAGCTCGTGTTG |
| Trial 1, colony 3 | | AGTAGCTTGCTACTTTGCTGACGAGCGGCGGACGGGTGAGTAATGCCTGGGAATATGCCTTGATGTGGGGGATAACTATTGGAAACGATAGCTAATACCGCATAATGTCTTCGGACCAAAGAGGGGGATCTTCGGACCTCTCGCGTCAAGATTAGCCCAGGTGAGATTAGCTAGTTGGTGGGGTAAGAGCTCACCAAGGCGACGATCTCTAGCTGGTCTGAGAGGATGATCAGCCACACTGGAACTGAGACACGGTCCAGACTCCTACGGGAGGCAGCAGTGGGGAATATTGCACAATGGGCGAAAGCCTGATGCAGCCATGCCGCGTGTATGAAGAAGGCCTTCGGGTTGTAAAGTACTTTCAGTCGTGAGGAAGGGTGTGCAGTTAATAGCTGCATATCTTGACGTTAGCGACAGAAGAAGCACCGGCTAACTCCGTGCCAGCAGCCGCGGTAATACGGAGGGTGCGAGCGTTAATCGGAATTACTGGGCGTAAAGCGCATGCAGGTGGTTCATTAAGTCAGATGTGAAAGCCCGGGGCTCAACCTCGGAACCGCATTTGAAACTGGTGAACTAGAGTGCTGTAGAGGGGGGTAGAATTTCAGGTGTAGCGGTGAAATGCGTAGAGATCTGAAGGAATACCAGTGGCGAAGGCGGCCCCCTGGACAGACACTGACACTCAGATGCGAAAGCGTGGGGAGCAAACAGGATTAGATACCCTGGTAGTCCACGCCGTAAACGATGTCTACTTGGAGGTTGTGGCCTTGAGCCGTGGCTTTCGGAGCTAACGCGTTAAGTAGACCGCCTGGGGAGTACGGTCGCAAGATTAAAACTCAAATGAATTGACGGGGGCCCGCACAAGCGGTGGAGCATGTGGTTTAATTCGATGCAACGCGAAGAACCTTACCTACTCTTGACATCTAC |
| Trial 1, colony 4 | | GTAGCTTGCTACTTTGCTGACGAGCGGCGGACGGGTGAGTAATGCCTGGGAATATGCCTTGATGTGGGGGATAACTATTGGAAACGATAGCTAATACCGCATAATGTCTTCGGACCAAAGAGGGGGATCTTCGGACCTCTCGCGTCAAGATTAGCCCAGGTGAGATTAGCTAGTTGGTGGGGTAAGAGCTCACCAAGGCGACGATCTCTAGCTGGTCTGAGAGGATGATCAGCCACACTGGAACTGAGACACGGTCCAGACTCCTACGGGAGGCAGCAGTGGGGAATATTGCACAATGGGCGAAAGCCTGATGCAGCCATGCCGCGTGTATGAAGAAGGCCTTCGGGTTGTAAAGTACTTTCAGTCGTGAGGAAGGGTGTGCAGTTAATAGCTGCATATCTTGACGTTAGCGACAGAAGAAGCACCGGCTAACTCCGTGCCAGCAGCCGCGGTAATACGGAGGGTGCGAGCGTTAATCGGAATTACTGGGCGTAAAGCGCATGCAGGTGGTTCATTAAGTCAGATGTGAAAGCCCGGGGCTCAACCTCGGAACCGCATTTGAAACTGGTGAACTAGAGTGCTGTAGAGGGGGGTAGAATTTCAGGTGTAGCGGTGAAATGCGTAGAGATCTGAAGGAATACCAGTGGCGAAGGCGGCCCCCTGGACAGACACTGACACTCAGATGCGAAAGCGTGGGGAGCAAACAGGATTAGATACCCTGGTAGTCCACGCCGTAAACGATGTCTACTTGGAGGTTGTGGCCTTGAGCCGTGGCTTTCGGAGCTAACGCGTTAAGTAGACCGCCTGGGGAGTACGGTCGCAAGATTAAAACTCAAATGAATTGACGGGGGCCCGCACAAGCGGTGGAGCATGTGGTTTAATTCGATGCAACGCGAAGAACCTTACCTACTCTTGACATCTA |
| Trial 1, colony 5 | | AGTAGCTTGCTACTTTGCTGACGAGCGGCGGACGGGTGAGTAATGCCTGGGAATATGCCTTGATGTGGGGGATAACTATTGGAAACGATAGCTAATACCGCATAATGTCTTCGGACCAAAGAGGGGGATCTTCGGACCTCTCGCGTCAAGATTAGCCCAGGTGAGATTAGCTAGTTGGTGGGGTAAGAGCTCACCAAGGCGACGATCTCTAGCTGGTCTGAGAGGATGATCAGCCACACTGGAACTGAGACACGGTCCAGACTCCTACGGGAGGCAGCAGTGGGGAATATTGCACAATGGGCGAAAGCCTGATGCAGCCATGCCGCGTGTATGAAGAAGGCCTTCGGGTTGTAAAGTACTTTCAGTCGTGAGGAAGGGTGTGCAGTTAATAGCTGCATATCTTGACGTTAGCGACAGAAGAAGCACCGGCTAACTCCGTGCCAGCAGCCGCGGTAATACGGAGGGTGCGAGCGTTAATCGGAATTACTGGGCGTAAAGCGCATGCAGGTGGTTCATTAAGTCAGATGTGAAAGCCCGGGGCTCAACCTCGGAACCGCATTTGAAACTGGTGAACTAGAGTGCTGTAGAGGGGGGTAGAATTTCAGGTGTAGCGGTGAAATGCGTAGAGATCTGAAGGAATACCAGTGGCGAAGGCGGCCCCCTGGACAGACACTGACACTCAGATGCGAAAGCGTGGGGAGCAAACAGGATTAGATACCCTGGTAGTCCACGCCGTAAACGATGTCTACTTGGAGGTTGTGGCCTTGAGCCGTGGCTTTCGGAGCTAACGCGTTAAGTAGACCGCCTGGGGAGTACGGTCGCAAGATTAAAACTCAAATGAATTGACGGGGGCCCGCACAAGCGGTGGAGCATGTGGTTTAATTCGATGCAACGCGAAGAACCTTACCTACTCTTGACATCTAC |
| Trial 1, colony 6 | | GTAGCTTGCTACTTTGCTGACGAGCGGCGGACGGGTGAGTAATGCCTGGGAATATGCCTTGATGTGGGGGATAACTATTGGAAACGATAGCTAATACCGCATAATGTCTTCGGACCAAAGAGGGGGATCTTCGGACCTCTCGCGTCAAGATTAGCCCAGGTGAGATTAGCTAGTTGGTGGGGTAAGAGCTCACCAAGGCGACGATCTCTAGCTGGTCTGAGAGGATGATCAGCCACACTGGAACTGAGACACGGTCCAGACTCCTACGGGAGGCAGCAGTGGGGAATATTGCACAATGGGCGAAAGCCTGATGCAGCCATGCCGCGTGTATGAAGAAGGCCTTCGGGTTGTAAAGTACTTTCAGTCGTGAGGAAGGGTGTGCAGTTAATAGCTGCATATCTTGACGTTAGCGACAGAAGAAGCACCGGCTAACTCCGTGCCAGCAGCCGCGGTAATACGGAGGGTGCGAGCGTTAATCGGAATTACTGGGCGTAAAGCGCATGCAGGTGGTTCATTAAGTCAGATGTGAAAGCCCGGGGCTCAACCTCGGAACCGCATTTGAAACTGGTGAACTAGAGTGCTGTAGAGGGGGGTAGAATTTCAGGTGTAGCGGTGAAATGCGTAGAGATCTGAAGGAATACCAGTGGCGAAGGCGGCCCCCTGGACAGACACTGACACTCAGATGCGAAAGCGTGGGGAGCAAACAGGATTAGATACCCTGGTAGTCCACGCCGTAAACGATGTCTACTTGGAGGTTGTGGCCTTGAGCCGTGGCTTTCGGAGCTAACGCGTTAAGTAGACCGCCTGGGGAGTACGGTCGCAAGATTAAAACTCAAATGAATTGACGGGGGCCCGCACAAGCGGTGGAGCATGTGGTTTAATTCGATGCAACGCGAAGAACCTTACCTACTCTTGACATCTACAGAATTCGCTAGAGATAGCTTAGTGCCTTCGGGAACTGTAAGACAGGTGCTGCATGGCTGTCGTCAGCTCGTGTTGTGAAATGTTGGGTTAAGTCCCGCAACGAGCGCAACCCTTATCCTTGTTTGCC |
| Trial 1, colony 7 | | AAGTAGCTTGCTACTTTGCTGACGAGCGGCGGACGGGTGAGTAATGCCTGGGAATATGCCTTGATGTGGGGGATAACTATTGGAAACGATAGCTAATACCGCATAATGTCTTCGGACCAAAGAGGGGGATCTTCGGACCTCTCGCGTCAAGATTAGCCCAGGTGAGATTAGCTAGTTGGTGGGGTAAGAGCTCACCAAGGCGACGATCTCTAGCTGGTCTGAGAGGATGATCAGCCACACTGGAACTGAGACACGGTCCAGACTCCTACGGGAGGCAGCAGTGGGGAATATTGCACAATGGGCGAAAGCCTGATGCAGCCATGCCGCGTGTATGAAGAAGGCCTTCGGGTTGTAAAGTACTTTCAGTCGTGAGGAAGGGTGTGCAGTTAATAGCTGCATATCTTGACGTTAGCGACAGAAGAAGCACCGGCTAACTCCGTGCCAGCAGCCGCGGTAATACGGAGGGTGCGAGCGTTAATCGGAATTACTGGGCGTAAAGCGCATGCAGGTGGTTCATTAAGTCAGATGTGAAAGCCCGGGGCTCAACCTCGGAACCGCATTTGAAACTGGTGAACTAGAGTGCTGTAGAGGGGGGTAGAATTTCAGGTGTAGCGGTGAAATGCGTAGAGATCTGAAGGAATACCAGTGGCGAAGGCGGCCCCCTGGACAGACACTGACACTCAGATGCGAAAGCGTGGGGAGCAAACAGGATTAGATACCCTGGTAGTCCACGCCGTAAACGATGTCTACTTGGAGGTTGTGGCCTTGAGCCGTGGCTTTCGGAGCTAACGCGTTAAGTAGACCGCCTGGGGAGTACGGTCGCAAGATTAAAACTCAAATGAATTGACGGGGGCCCGCACAAGCGGTGGAGCATGTGGTTTAATTCGATGCAACGCGAAGAACCTTACCTACTCTTGACATCTAC |
| Trial 1, colony 8 | | GCTGACGAGCGGCGGACGGGTGAGTAATGCCTGGGAATATGCCTTGATGTGGGGGATAACTATTGGAAACGATAGCTAATACCGCATAATGTCTTCGGACCAAAGAGGGGGATCTTCGGACCTCTCGCGTCAAGATTAGCCCAGGTGAGATTAGCTAGTTGGTGGGGTAAGAGCTCACCAAGGCGACGATCTCTAGCTGGTCTGAGAGGATGATCAGCCACACTGGAACTGAGACACGGTCCAGACTCCTACGGGAGGCAGCAGTGGGGAATATTGCACAATGGGCGAAAGCCTGATGCAGCCATGCCGCGTGTATGAAGAAGGCCTTCGGGTTGTAAAGTACTTTCAGTCGTGAGGAAGGGTGTGTAGTTAATAGCTGCATATCTTGACGTTAGCGACAGAAGAAGCACCGGCTAACTCCGTGCCAGCAGCCGCGGTAATACGGAGGGTGCGAGCGTTAATCGGAATTACTGGGCGTAAAGCGCATGCAGGTGGTTCATTAAGTCAGATGTGAAAGCCCGGGGCTCAACCTCGGAACCGCATTTGAAACTGGTGAACTAGAGTGCTGTAGAGGGGGGTAGAATTTCAGGTGTAGCGGTGAAATGCGTAGAGATCTGAAGGAATACCAGTGGCGAAGGCGGCCCCCTGGACAGACACTGACACTCAGATGCGAAAGCGTGGGGAGCAAACAGGATTAGATACCCTGGTAGTCCACGCCGTAAACGATGTCTACTTGGAGGTTGTGGCCTTGAGCCGTGGCTTTCGGAGCTAACGCGTTAAGTAGACCGCCTGGGGAGTACGGTCGCAAGATTAAAACTCAAATGAATTGACGGGGGCCCGCACAAGCGGTGGAGCATGTGGTTTAATTCGATGCAACGCGAAGAACCTTACCTACTCTTGACATCTACAGAATTCGCTAGAGATAGCTTAGTGCCTTCGGGAACTGTAAGACAGGTGCTGCATGGCTGTCGTCAGCTCGTGTTGTGAAATGTTGG |
| Trial 1, colony 9 | | GTAGCTTGCTACTTTGCTGACGAGCGGCGGACGGGTGAGTAATGCCTGGGAATATGCCTTGATGTGGGGGATAACTATTGGAAACGATAGCTAATACCGCATAATGTCTTCGGACCAAAGAGGGGGATCTTCGGACCTCTCGCGTCAAGATTAGCCCAGGTGAGATTAGCTAGTTGGTGGGGTAAGAGCTCACCAAGGCGACGATCTCTAGCTGGTCTGAGAGGATGATCAGCCACACTGGAACTGAGACACGGTCCAGACTCCTACGGGAGGCAGCAGTGGGGAATATTGCACAATGGGCGAAAGCCTGATGCAGCCATGCCGCGTGTATGAAGAAGGCCTTCGGGTTGTAAAGTACTTTCAGTCGTGAGGAAGGGTGTGCAGTTAATAGCTGCATATCTTGACGTTAGCGACAGAAGAAGCACCGGCTAACTCCGTGCCAGCAGCCGCGGTAATACGGAGGGTGCGAGCGTTAATCGGAATTACTGGGCGTAAAGCGCATGCAGGTGGTTCATTAAGTCAGATGTGAAAGCCCGGGGCTCAACCTCGGAACCGCATTTGAAACTGGTGAACTAGAGTGCTGTAGAGGGGGGTAGAATTTCAGGTGTAGCGGTGAAATGCGTAGAGATCTGAAGGAATACCAGTGGCGAAGGCGGCCCCCTGGACAGACACTGACACTCAGATGCGAAAGCGTGGGGAGCAAACAGGATTAGATACCCTGGTAGTCCACGCCGTAAACGATGTCTACTTGGAGGTTGTGGCCTTGAGCCGTGGCTTTCGGAGCTAACGCGTTAAGTAGACCGCCTGGGGAGTACGGTCGCAAGATTAAAACTCA |
| Trial 1, colony 10 | | AAGTAGCTTGCTACTTTGCTGACGAGCGGCGGACGGGTGAGTAATGCCTGGGAATATGCCTTGATGTGGGGGATAACTATTGGAAACGATAGCTAATACCGCATAATGTCTTCGGACCAAAGAGGGGGATCTTCGGACCTCTCGCGTCAAGATTAGCCCAGGTGAGATTAGCTAGTTGGTGGGGTAAGAGCTCACCAAGGCGACGATCTCTAGCTGGTCTGAGAGGATGATCAGCCACACTGGAACTGAGACACGGTCCAGACTCCTACGGGAGGCAGCAGTGGGGAATATTGCACAATGGGCGAAAGCCTGATGCAGCCATGCCGCGTGTATGAAGAAGGCCTTCGGGTTGTAAAGTACTTTCAGTCGTGAGGAAGGGTGTGCAGTTAATAGCTGCATATCTTGACGTTAGCGACAGAAGAAGCACCGGCTAACTCCGTGCCAGCAGCCGCGGTAATACGGAGGGTGCGAGCGTTAATCGGAATTACTGGGCGTAAAGCGCATGCAGGTGGTTCATTAAGTCAGATGTGAAAGCCCGGGGCTCAACCTCGGAACCGCATTTGAAACTGGTGAACTAGAGTGCTGTAGAGGGGGGTAGAATTTCAGGTGTAGCGGTGAAATGCGTAGAGATCTGAAGGAATACCAGTGGCGAAGGCGGCCCCCTGGACAGACACTGACACTCAGATGCGAAAGCGTGGGGAGCAAACAGGATTAGATACCCTGGTAGTCCACGCCGTAAACGATGTCTACTTGGAGGTTGTGGCCTTGAGCCGTGGCTTTCGGAGCTAACGCGTTAAGTAGACCGCCTGGGGAGTACGGTCGCAAGATTAAAACTCAAATGAATTGACGGGGGCCCGCACAAGCGGTGGAGCATGTGGTTTAATTCGATGCAACGCGAAGAACCTTACCTACTCTTGACATCTACAGAATTCGCTAGAGATAGCTTAGTGCCTTCGGGAACTGTAAGACAGGTGCTGCATGGCTGTCG |
| Trial 2.2, colony 1 | | GACGAGCGGCGGACGGGTGAGTAATGCCTGGGAATATGCCTTGATGTGGGGGATAACTATTGGAAACGATAGCTAATACCGCATAATGTCTTCGGACCAAAGAGGGGGATCTTCGGACCTCTCGCGTCAAGATTAGCCCAGGTGAGATTAGCTAGTTGGTGGGGTAAGAGCTCACCAAGGCGACGATCTCTAGCTGGTCTGAGAGGATGATCAGCCACACTGGAACTGAGACACGGTCCAGACTCCTACGGGAGGCAGCAGTGGGGAATATTGCACAATGGGCGAAAGCCTGATGCAGCCATGCCGCGTGTATGAAGAAGGCCTTCGGGTTGTAAAGTACTTTCAGTCGTGAGGAAGGGTGTGTAGTTAATAGCTGCATATCTTGACGTTAGCGACAGAAGAAGCACCGGCTAACTCCGTGCCAGCAGCCGCGGTAATACGGAGGGTGCGAGCGTTAATCGGAATTACTGGGCGTAAAGCGCATGCAGGTGGTTCATTAAGTCAGATGTGAAAGCCCGGGGCTCAACCTCGGAACCGCATTTGAAACTGGTGAACTAGAGTGCTGTAGAGGGGGGTAGAATTTCAGGTGTAGCGGTGAAATGCGTAGAGATCTGAAGGAATACCAGTGGCGAAGGCGGCCCCCTGGACAGACACTGACACTCAGATGCGAAAGCGTGGGGAGCAAACAGGATTAGATACCCTGGTAGTCCACGCCGTAAACGATGTCTACTTGGAGGTTGTGGCCTTGAGCCGTGGCTTTCGGAGCTAACGCGTTAAGTAGACCGCCTGGGGAGTACGGTCGCAAGATTAAAACTCAAATGAATTGACGGGGGCCCGCACAAGCGGTGGAGCATGTGGTTTAATTCGATGCAACGCGAAGAACCTTACCTACTCTTGACATC |
| Trial 2.2, colony 2 | | GCTTGCTAATTTGCTGACGAGCGGCGGACGGGTGAGTAATGCCTGGGAATATGCCTTGATGTGGGGGATAACTATTGGAAACGATAGCTAATACCGCATAATGTCTTCGGACCAAAGAGGGGGATCTTCGGACCTCTCGCGTCAAGATTAGCCCAGGTGAGATTAGCTAGTTGGTGGGGTAAGAGCTCACCAAGGCGACGATCTCTAGCTGGTCTGAGAGGATGATCAGCCACACTGGAACTGAGACACGGTCCAGACTCCTACGGGAGGCAGCAGTGGGGAATATTGCACAATGGGCGAAAGCCTGATGCAGCCATGCCGCGTGTATGAAGAAGGCCTTCGGGTTGTAAAGTACTTTCAGTCGTGAGGAAGGGTGTGTAGTTAATAGCTGCATATCTTGACGTTAGCGACAGAAGAAGCACCGGCTAACTCCGTGCCAGCAGCCGCGGTAATACGGAGGGTGCGAGCGTTAATCGGAATTACTGGGCGTAAAGCGCATGCAGGTGGTTCATTAAGTCAGATGTGAAAGCCCGGGGCTCAACCTCGGAACCGCATTTGAAACTGGTGAACTAGAGTGCTGTAGAGGGGGGTAGAATTTCAGGTGTAGCGGTGAAATGCGTAGAGATCTGAAGGAATACCAGTGGCGAAGGCGGCCCCCTGGACAGACACTGACACTCAGATGCGAAAGCGTGGGGAGCAAACAGGATTAGATACCCTGGTAGTCCACGCCGTAAACGATGTCTACTTGGAGGTTGTGGCCTTGAGCCGTGGCTTTCGGAGCTAACGCGTTAAGTAGACCGCCTGGGGAGTACGGTCGCAAGATTAAAACTCAAATGAATTGACGGGGGCCCGCACAAGCGGTGGAGCATGTGGTTTAATTCGATGCAACGCGAAGAACCTTACCTACTCTTGACATCTACAGAATTCGCTAGAGATAGCTTAGTGCCTTCGGGAACTGTAAGACAGGTGCTGCATGGCTGTCGTCAGCTCGTGTTGTGAAATGTTGGGTTAAGTCCCGCAACGAGCGCAACCCTTATCCTT |
